# Pyrfume: A Window to the World’s Olfactory Data

**DOI:** 10.1101/2022.09.08.507170

**Authors:** Jason B. Castro, Travis J. Gould, Robert Pellegrino, Zhiwei Liang, Liyah A. Coleman, Famesh Patel, Derek S. Wallace, Tanushri Bhatnagar, Joel D. Mainland, Richard C. Gerkin

## Abstract

Advances in theoretical understanding are frequently unlocked by access to large, diverse experimental datasets. Olfactory neuroscience and psychophysics remain years behind the other senses in part because rich datasets linking olfactory stimuli with their corresponding percepts, behaviors, and neural pathways underlying this transformation, remain scarce. Here we present a concerted effort to unlock and unify dozens of stimulus-linked olfactory datasets across species and modalities under a unified framework called *Pyrfume*. We present examples of how researchers might use Pyrfume to conduct novel analyses uncovering new principles, introduce trainees to the field, or construct benchmarks for machine olfaction.

## INTRODUCTION

Mapping from the high dimensional space of chemical features to the much lower dimensional spaces of percepts, decisions, and behaviors is the quintessential feat pulled off by the chemical senses^1–4^. However, despite considerable effort, we understand this mapping only very poorly, and in limited special cases^5,6^. Missing theories are partly to blame, but an equally salient and fundamental problem is missing data. Simply put, the data have not existed to answer--or even adequately ask--the most central questions in the field. We hypothesize that with the right data, ‘solving’ olfaction will ultimately prove conceptually tractable, much like the sensory phenomena of pitch perception or color vision – two success stories of classical psychophysics that were well underway in the 19th century^7,8^.

Fundamental coding and perceptual principles for pitch and color vision can be studied in experiments where simple stimuli are both varied smoothly and manipulated easily. This is not the case in olfaction; to the contrary, olfactory stimulus space is massive^9^, laborious to explore, and difficult to digitize. No one investigator in any one study has a realistic hope of probing a large portion of odor space (e.g. thousands of odorants) in conjunction with the full gamut of sensory or neurophysiological measurements. However, if one could make effective use of the space of all investigators, and pool data across many studies, many fundamental questions may actually already be within striking distance. With the right data and pipeline for standardization, olfaction can be made much more amenable to the machine learning approaches that have proven so fruitful and generative in other sensory modalities.

The tremendous success of data-driven approaches for problems in visual coding and scene analysis have been driven by the wide availability and accessibility of imaging data, as well as the communal adoption of key datasets for testing and benchmarking^6,10,11^. In the computer vision community, for example, the MNIST^12^ and ImageNet^13^ datasets are understood to be essential proving grounds for any newly proposed algorithm or coding principle. Additionally, new visual coding theories can be quickly tested and prototyped across a large number of heterogeneous datasets that expose them to different contexts and edge-cases, making models more robust through out-of-sample testing^14^. Here, we describe a newly curated set of >40 olfactory datasets and a new suite of data fetching, management, and curation tools that we believe can create an “MNIST for olfaction”, and help stimulate a new era of data-driven inquiry in the olfaction community.

In the remainder of this paper, we introduce and demonstrate the functionality of *Pyrfume*, an integrated data archive with companion python library, and python, R, and REST APIs for accelerating inquiry in olfactory neuroscience. While the importance of data aggregation in olfaction has been recognized by others, and there have been other efforts on this front, Pyrfume is notable for its breadth and coverage, spanning >40 odorant-linked datasets in mammalian olfaction including human psychophysics and perception; and mammalian psychophysics, behavior, brain imaging, physiology, and pharmacology. All together, it contains information about >20,000 identified odorants. We believe that Pyrfume will prove useful in facilitating communication between two research communities who have much to learn from each other, but who have interacted only modestly in the past: 1) basic scientists studying olfaction, who would like their data to be among the standard datasets used in tasks centered on modeling and prediction, and 2) machine learning researchers with a theoretical interest in olfaction, but who have not known how to quickly find the relevant data.

### CORE DESIGN PRINCIPLES OF PYRFUME

There are many archives, papers, and search engines with data that are useful to olfactory scientists, but it has been impossible to coordinate structured queries across these, and each alone has limitations pertaining to size, coverage, or accessibility. PubChem^15^, for example, has detailed chemical information for over 10M molecules, but has very little olfactory data. The well-studied Dravnieks atlas^16^ data set has molecules whose perceptual qualities are described with a structured vocabulary amenable to machine learning, but contains data for only 138 unique molecules. The National Geographic Smell Survey^17^ has odor intensity ratings collected from an impressive 1.4M subjects, but only for a total of 6 odorants. Even for investigators with the motivation and technical acumen to integrate olfactory data from different sources, there are still the additional challenges of scraping, cleaning, and “munging” datasets that are effectively siloed in separate repositories, and which employ idiosyncratic formats for organizing data.

Pyrfume aims to overcome these limitations, and is premised on the simple ideas that: 1) most olfaction experiments are straightforward to index (on the unique identifier (ID) of an olfactory stimulus, e.g. molecules, substances, or mixtures at a given concentration), and 2) any olfactory experiment can be generically described as a machine-readable pairing of such stimulus IDs, the task performed with the corresponding stimuli, and the observed individuals and their behaviors. Linking these experimental components allows for a robust data-formatting standard which is flexible enough to accommodate a wide array of experimental designs and data types, and which conforms to principles of good database design. Note that *behavior*, as used here and throughout the paper, refers to any experimental measurement. These could be human perceptual ratings applied to a given chemical, glomerular responses observed in mouse physiology experiments, or measured sensitivities of receptors in pharmacology experiments, etc. (**Figure 1**).

**Figure 1.**
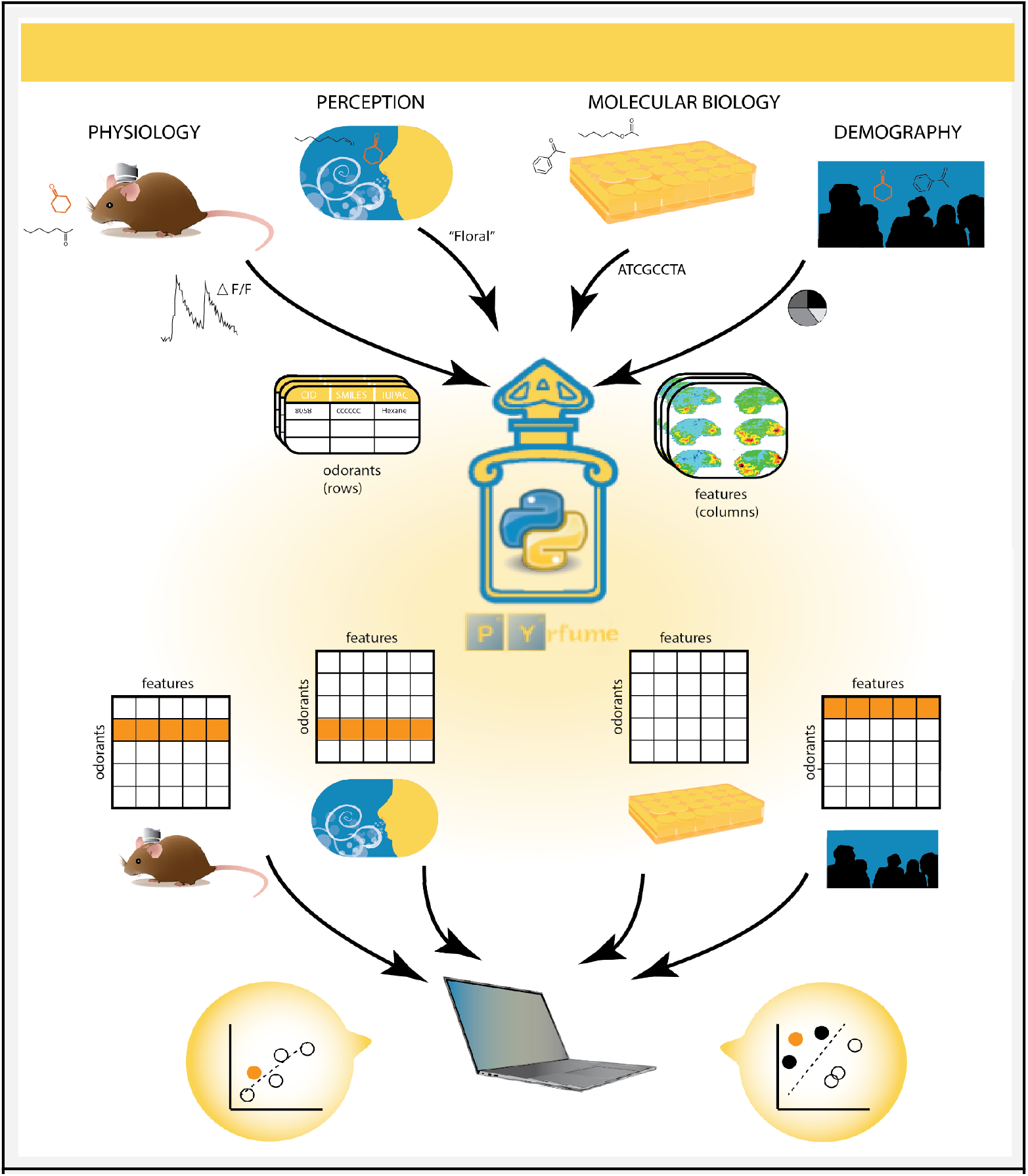
Overview of the Pyrfume ecosystem. Disparate and heterogeneous data are ‘bottled’ (top) to make them amenable to cross-modal, meta-analysis, or machine learning (ML) tasks, which typically require tabular data in the form of samples x features matrices. Under the Pyrfume standard, data are always linked to the odorant stimuli comprising a given experiment. The features will depend on the particular experiment, but could include data such as amplitudes of glomerular calcium transients, vectorized perceptual descriptors, receptor sensitivities, etc. Bottled experiments are publicly available through Python, R, or REST APIs or directly via GitHub. Any given data archive on Pyrfume (>40 to date) can be easily fetched, or ‘unbottled’ by a user using the Pyrfume API, and used immediately for ML tasks (bottom), with no need for laborious cleaning or formatting. The orange rows of the odorant x feature matrices indicate odorant molecules common to the experiments. The ability to easily extract data about common odorants across experimental modalities and model systems is a unique strength of Pyrfume.

Each data source curated in Pyrfume is standardized to conform to a subject/object design framework, separating the odor objects from behavior of the subject(s) under study. At the object level, the most essential file, called `stimuli.csv` is indexed on a stimulus ID and maps this ID to the chemical/molecular details of the odorants used. A stimulus could represent a single molecule, substance, or mixture, the applied concentration(s), and potentially other experimental conditions. In (typical) cases where at least one stimulus is a single molecule with known structure, the Pyrfume archive will also contain a file, `molecules.csv`, that lists all molecules used in that dataset, with columns providing PubChem Compound IDs (CIDs), SMILES^18^, common names, and IUPAC names. This file is useful for indexing the usage of each kind of molecule across datasets, and also for computing physicochemical features for each such molecule (software packages such as RDKit and Mordred^19^ compute these directly from a SMILES). When stimuli correspond to mixtures of unknown provenance, e.g. “cloves”, a unique stimulus ID is generated but in such cases it may not be possible to link it back to specific compounds in `molecules.csv`. In cases where a dataset includes calculated or experimentally measured physicochemical properties of molecules, these may be included in `physics.csv`, typically indexed on CID. Collectively these describe the odorant “object”.

The subject side of the data is principally described in `behavior.csv`, a (typically) a long-format dataframe, indexed on stimulus ID. In a simple (hypothetical) experiment where, say, 100 molecules were each rated on 2 perceptual qualities (‘sweetness’, ‘floral-ness’), the behavior file would have dimensions 200 × 3, with an indexing column for perceptual quality:

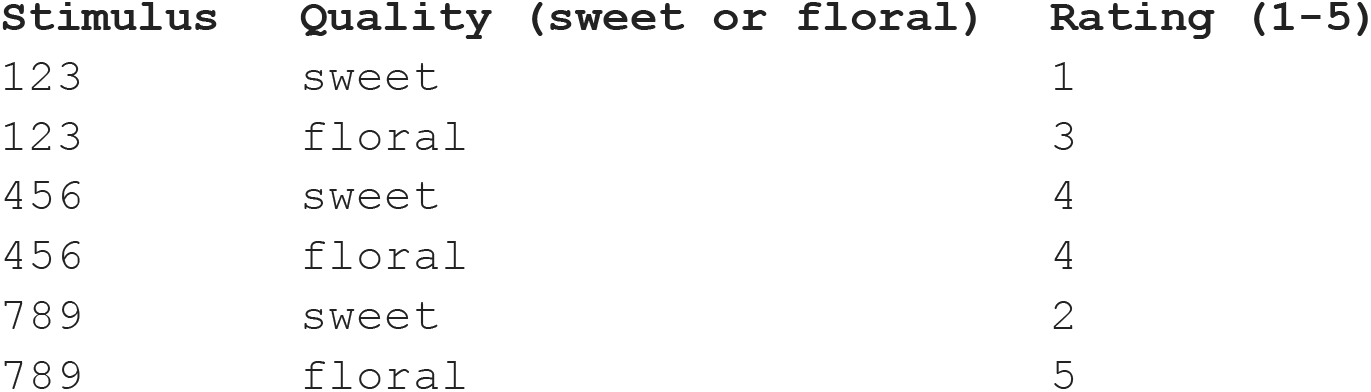

This standard, widely referred to as ‘Tidy’, or sometimes ‘Third normal form’^20^, can easily accommodate more complex, multi-level designs, and reduces both redundancy in data representation, as well as the unnecessary proliferation of files and tables. In the Pyrfume archive that corresponds to the glomerular imaging data of Chae et al., 2019^21^, for example, there are 5 animals, 2 hemibulbs, both high and low concentration experimental conditions, and varying numbers of imaged glomeruli in each experiment. Conversion to formats that feel more natural for many experimentalists (i.e. tables of odorants x glomerular responses, with one table per condition) is accomplished easily with standard aggregation and pivoting steps, 1-2 lines of code in Python (via Pandas^22^) or in R.

In cases where stimuli are administered to distinct subjects, a `subjects.csv` file provides a mapping between a subject ID and that subject’s distinguishing features. A ‘subject’ could be a human (e.g. identified by age, gender, region, etc.) participating in an odor perception study or a glomerulus (e.g. identified by mouse, olfactory bulb hemisphere, etc.) being studied in a physiological experiment. Collectively `subjects.csv` and `behavior.csv` describe the “subject” of the experiment and its behavior(s). These files are depicted in **Figure 2**.

**Figure 2.**
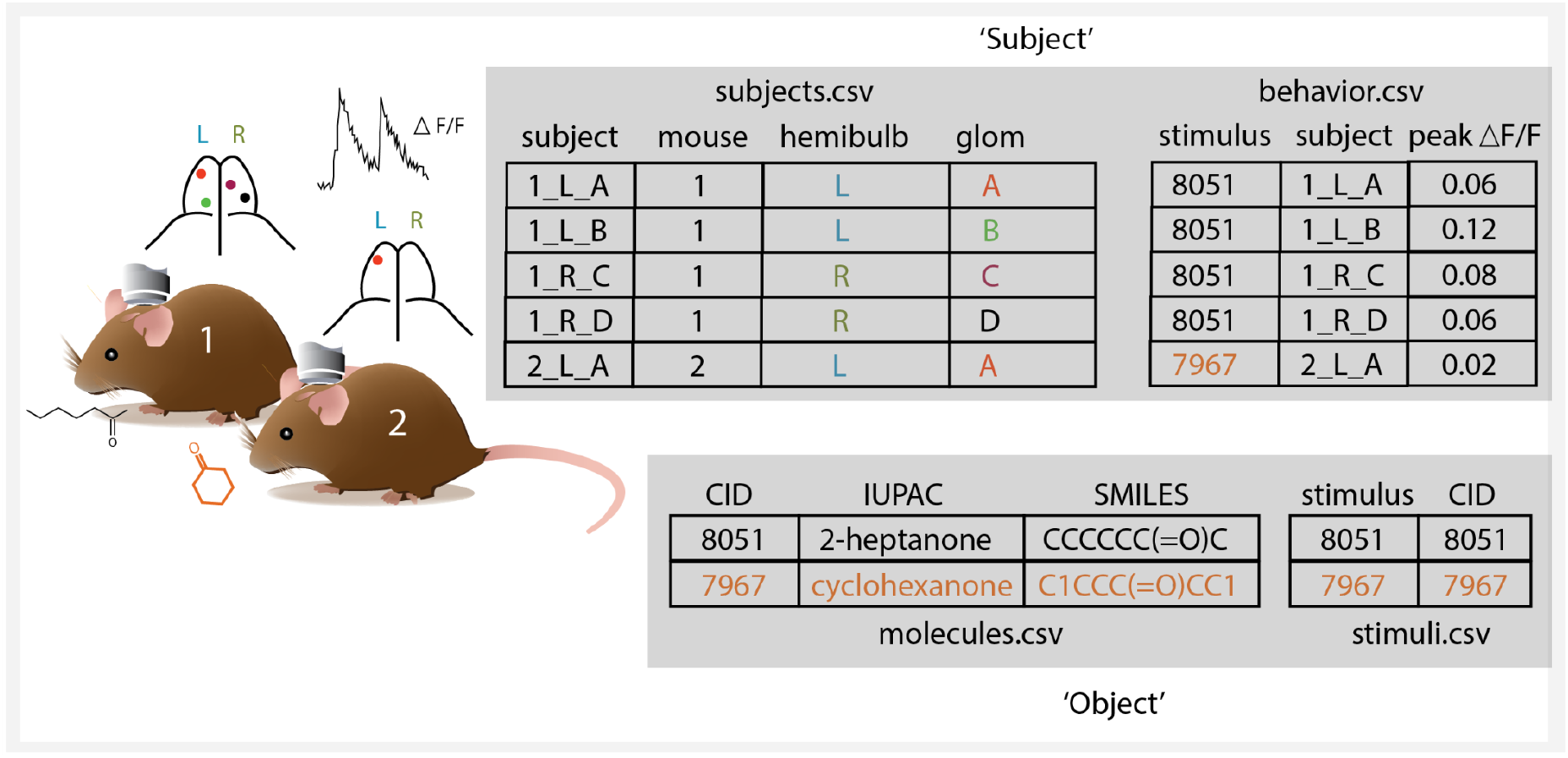
Schematic showing important data-formatting standards of Pyrfume. In the hypothetical and simplified experiment here, 2 odorants were tested on two animals, and data were collected across a total of 5 glomeruli.. The Pyrfume archive is organized around a subject-object distinction. The molecules.csv file (lower left) contains chemical descriptors, which can be used to index experimental measurements, and from which molecular features can be computed using packages such as RDKit^23^ or Mordred^19^. The stimuli.csv file – which is somewhat trivial in this case, for ease of illustration – provides a mapping between stimulus conditions and chemical information. The subjects.csv file (upper left) catalogs all experimental levels for all subjects, in ‘tidy’ form^20^. Lastly, behavior.csv is indexed on the stimuli, and contains the actual experimental measurements, with one measurement per row.

Lastly, each dataset archived includes two additional files: 1) `main.py`, a Python script of the data processing workflow, and 2) a simple, standardized, machine-readable markup file called `manifest.toml`, which describes the data archive. Using the TOML file format^24^, archive manifests contain sections to cite and credit relevant sources (`[source]`), list raw (`[raw]`) and processed (`[processed]`) data files included in the archive while also providing any important context or metadata for interpreting the data, and lists of any code used to obtain or process raw data ([code]). In sum, a typical complete, archived object in Pyrfume consists of at up to 7 files: `manifest.toml`, `main.py`, `stimuli.csv`, `molecules.csv`, `behavior.csv`, `subjects.csv`, and `physics.csv`, though the latter two are found only where applicable.

Uniquely identifying the stimuli--in particular correctly resolving them to precise molecular structures--is necessary for linking across datasets and for testing predictive models. In most cases this is achieved by first determining the PubChem CID--a unique structural identifier--corresponding to molecules described in the raw data. The pyrfume `get_cids()` function achieves this purpose. The CID can then be used to programmatically generate alternative identifiers using the PubChem API, through the pyrfume function `from_cids()`. In rare cases where a molecule does not correspond to any compound indexed in PubChem, an alternative identifier can be used directly. The decision to use PubChem CIDs for indexing molecules is motivated by the strength of PubChem as a source for integrating heterogeneous information about compounds (including safety, availability, and physical properties), and because SMILES lack idempotency. InChiKey would have been a solid alternative choice.

All examples here show usage of the Python API (http://github.com/pyrfume/pyrfume), which uses a local cache of data to avoid re-downloading the same data more than once. The name *pyrfume* follows the whimsical pythonic naming tradition of inserting the letters *py* into a word describing the subject matter (with perfume being an example of an odorous substance). A subset of the functionality shown here is also available through an R API (http://github.com/pyrfume/rfume). Both utilize a REST API to obtain data from the remote archives on GitHub, and this REST API can be used directly for those who wish to build additional tools in other programming languages.

### SAMPLE USE CASES

The basic Pyrfume ecosystem is described in **Figure 1**, and is defined by two ways of interacting with data, which we call *bottling* and *unbottling*. Bottling involves investigators readying their experimental data for machine learning applications, and the goal of this process is essentially to make life easy for downstream users (data scientists, computational neuroscientists, machine learning engineers, etc.) through standardization. In a typical workflow, an investigator would compile an inventory of all odorants used in their experiments, and then use the Pyrfume functions `get_cids()` and `from_cids()` to programatically create the `molecules.csv` file. **Table 1** lists and describes those pyrfume functions (of ∼70 total), that are most frequently used by a typical user. Creation of the `behavior.csv` file is more idiosyncratic to the particular experiment under consideration, but is rarely more than an hour’s work. The critical step, as alluded to above, involves defining the measurements that will comprise individual cell values of the dataframe (e.g. “each cell = peak deltaF/F measured in one glomerulus, for one odorant”, or “each cell = an EC50 value reported for one receptor, for one odorant”, or “each cell = a perceptual rating applied for one descriptor, for one odorant”, etc). As of the writing of this manuscript, there are >40 bottled experiments, comprising data for >20,000 unique odorants. In addition to data assembled from supplemental materials of published research, Pyrfume has cleaned and digitized versions of several large databases, which are shown in **Table 2**.

**Table 1.** The most commonly used Pyrfume functions for creating and working with archives.

| Function | Module | Arguments | Returns |
| --- | --- | --- | --- |
| `list_archives()` | base |  | List of Pyrfume archives (datasets) |
| `show_files()` | base | archive name | List of archive files in manifest |
| `load_manifest()` | base | archive name | Contents of archive manifest |
| `load_data()` | base | relative path to archive data file | Contents of data file |
| `get_cid()` | odorants | identifier (CAS, name, SMILES, etc) | PubChem Compound ID (CID) |
| `get_cids()` | odorants | list of identifiers | Corresponding list of CIDs |
| `from_cids()` | odorants | list of CIDs | List of dictionaries containing CID, Molecular Weight, IsomericSMILES, IUPAC name, and common name for each compound |

**Table 2.**
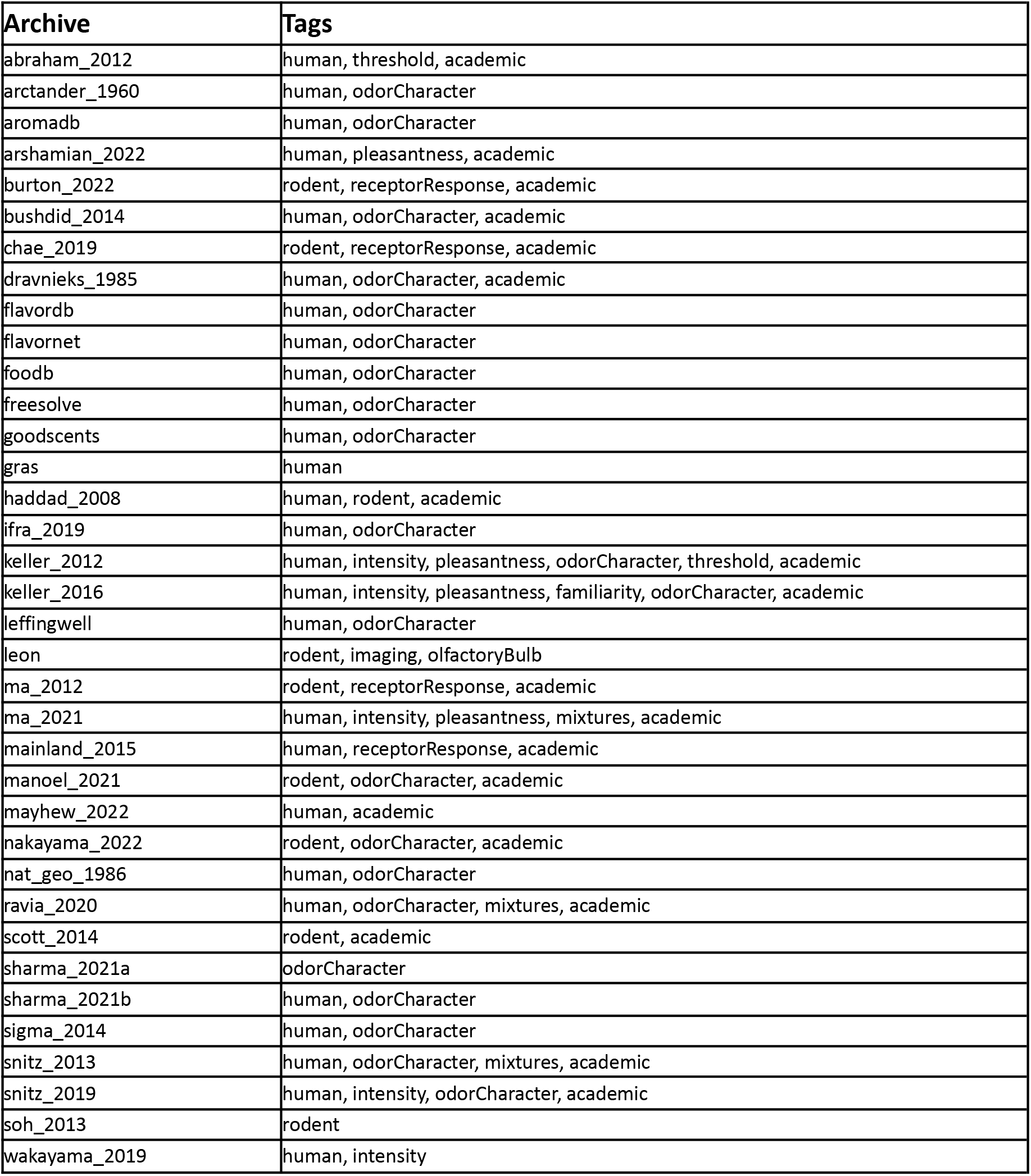
Inventory of data currently available through Pyrfume. Data currently available in standardized form through Pyrfume, as shown via the Python API (`pyrfume.list_archives()`). This can alternatively be viewed directly on the GitHub landing page.

| Archive | Tags |
| --- | --- |
| abraham_2012 | human, threshold, academic |
| arctander_1960 | human, odorCharacter |
| aromadb | human, odorCharacter |
| arshamian_2022 | human, pleasantness, academic |
| burton_2022 | rodent, receptorResponse, academic |
| bushdid_2014 | human, odorCharacter, academic |
| chae_2019 | rodent, receptorResponse, academic |
| dravnieks_1985 | human, odorCharacter, academic |
| flavordb | human, odorCharacter |
| flavornet | human, odorCharacter |
| foodb | human, odorCharacter |
| freesolve | human, odorCharacter |
| goodscents | human, odorCharacter |
| gras | human |
| haddad_2008 | human, rodent, academic |
| ifra_2019 | human, odorCharacter |
| keller_2012 | human, intensity, pleasantness, odorCharacter, threshold, academic |
| keller_2016 | human, intensity, pleasantness, familiarity, odorCharacter, academic |
| leffingwell | human, odorCharacter |
| leon | rodent, imaging, olfactoryBulb |
| ma_2012 | rodent, receptorResponse, academic |
| ma_2021 | human, intensity, pleasantness, mixtures, academic |
| mainland_2015 | human, receptorResponse, academic |
| manoel_2021 | rodent, odorCharacter, academic |
| mayhew_2022 | human, academic |
| nakayama_2022 | rodent, odorCharacter, academic |
| nat_geo_1986 | human, odorCharacter |
| ravia_2020 | human, odorCharacter, mixtures, academic |
| scott_2014 | rodent, academic |
| sharma_2021a | odorCharacter |
| sharma_2021b | human, odorCharacter |
| sigma_2014 | human, odorCharacter |
| snitz_2013 | human, odorCharacter, mixtures, academic |
| snitz_2019 | human, intensity, odorCharacter, academic |
| soh_2013 | rodent |
| wakayama_2019 | human, intensity |

### CROSS-MODAL ANALYSIS

For a concrete example of using Pyrfume, suppose an investigator were interested in the similarities between human and mouse odor representations. Some level of homology is presumed in any study where coding principles in one species are generalized to another, but this is actually an important empirical question that has been infrequently addressed^4,25,26^. A natural question is: do categories of human odor percepts correspond to categories of odor-evoked neuronal responses in mice? The data exist for addressing this, but without Pyrfume, these kinds of queries across datasets and species require the laborious tracking down of relevant studies, inventorying of chemicals used in each study, and cleaning and formatting of each study’s data so that direct comparisons are meaningful. With Pyrfume, this is all handled with 3 lines of code.

To conduct this inquiry much more straightforwardly with Pyrfume (**Figure 3**), an investigator or data analyst would scan the current contents of the database using the Pyrfume data tracker (https://status.pyrfume.org), and filter those datasets with the tags `human`, `odor character`, and `rodent`. For purposes of illustration, we will suppose that the datasets of Dravnieks et al, 1985^16^ (human odor qualities), and Chae et al, 2019^21^ (mouse glomerular imaging) were further identified to be of particular interest. An investigator would open any Python environment in which Pyrfume has been installed, and simply run the commands:

**Figure 3.**
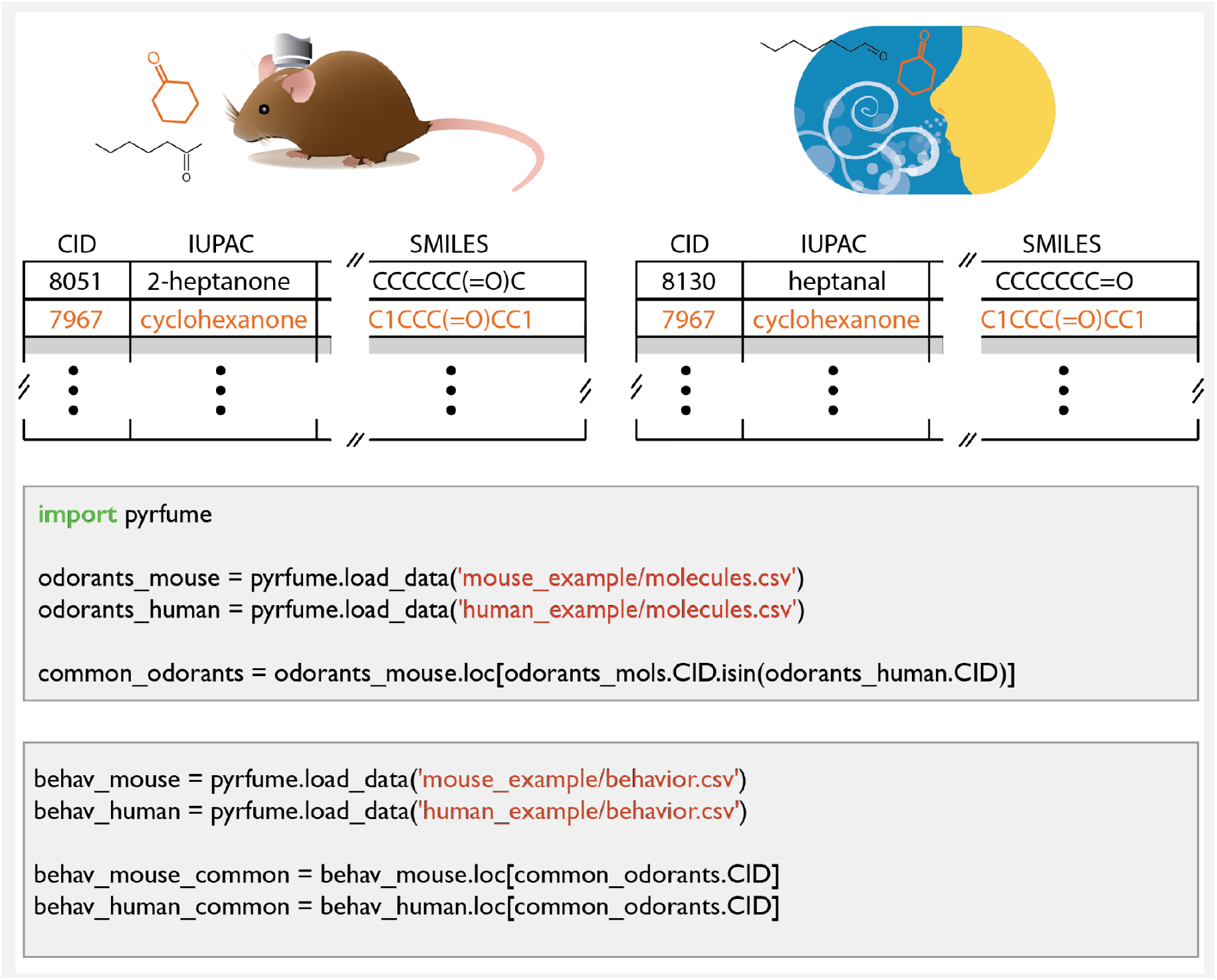
Illustration of methods for cross-experiment comparisons. Top: Molecule dataframes for hypothetical mouse and human experiments, which may have odorants in common (indicated in orange). The gray cells below show sample python code for extracting common odorants, and extracting subsets of data with shared odorants. The code shown is syntactically correct, and accurately represents the complexity (or simplicity) of these tasks, but the data sets are hypothetical for purposes of illustration.

~~~
molecules_dravnieks = pyrfume.get_data(dravnieks_1985/molecules.csv)
behaviors_dravnieks = pyrfume.get_data(dravnieks_1985/behavior_1.csv)
molecules_chae = pyrfume.get_data(chae_2019/molecules.csv)
behaviors_chae = pyrfume.get_data(chae_2019/behavior_1.csv)
~~~

Where the “_1” suffix distinguishes the specific behavior/task investigated here from other behaviors/tasks curated from the same dataset (“_2”, “_3”, etc., as described in the corresponding `manifest.toml` files or via `load_manifest()`). The objects returned require no additional cleaning, and are immediately ready for machine learning tasks using standard packages such as pandas and scikit-learn^27^. Obtaining molecules common to both the mouse and human datasets is achieved through simple, standard boolean and set operations in pandas.

As an illustration of a simple analysis, we performed spectral bi-clustering^28^ on an odorants x qualities matrix obtained in one step from `behaviors_dravnieks`, for those shared odorants contained in both `molecules_dravnieks` and `molecules_chae`. Biclustering is essentially a permutation of rows and columns that reveals any underlying block, or ‘checkerboard’ structure to a matrix. The clustered Dravnieks data clearly show three classes of molecules, characterized by unique sets of perceptual descriptors (**Figure 4**). Permuting the rows of the Chae glomerular data using the same row indices reveals that glomerular patterns also tend to fall into three classes that track the perceptual groups.

**Figure 4.**
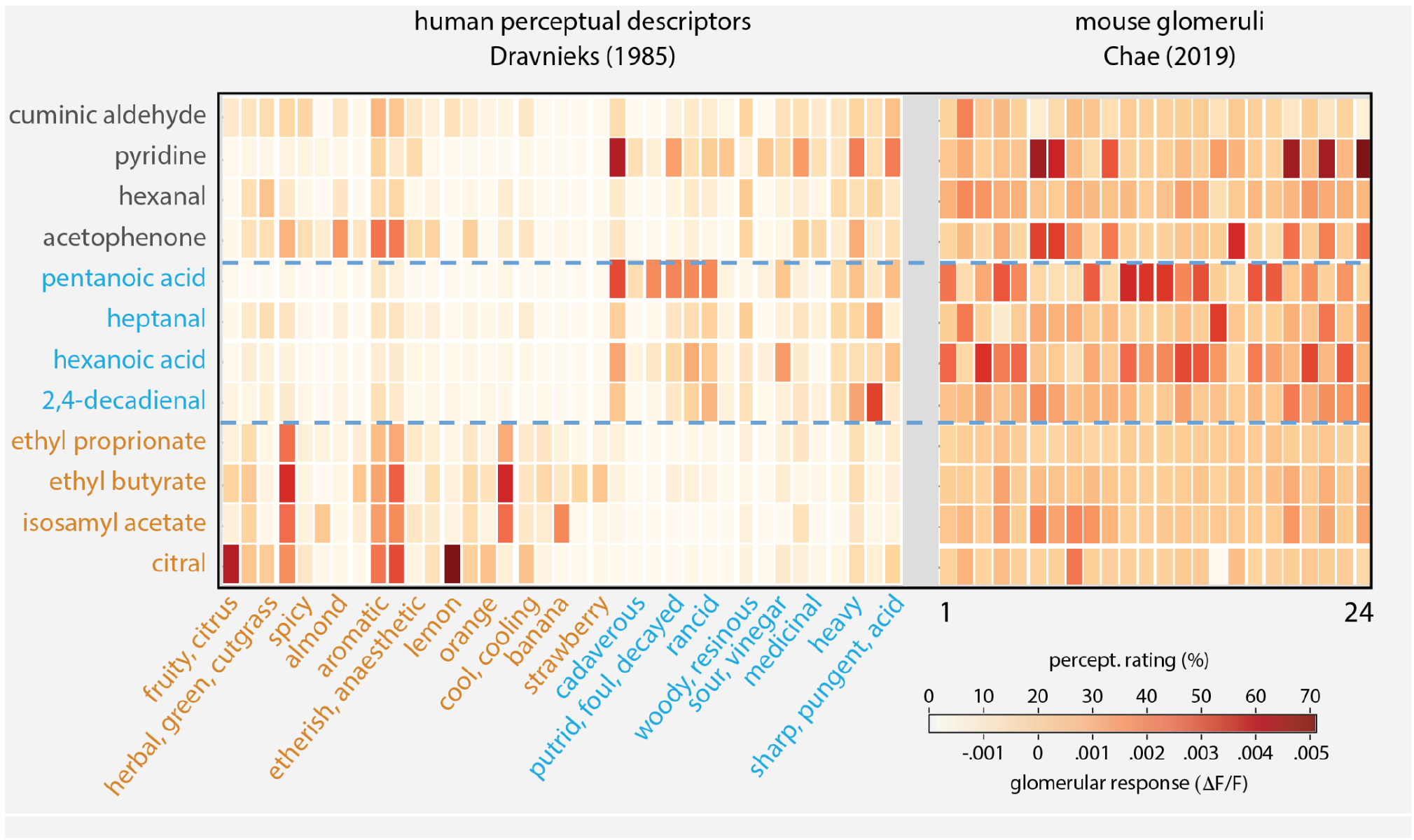
Human perceptual descriptions and mouse glomerular responses cluster similarly. Data from Dravnieks, 1985^16^ were clustered, for those odorants in common with Chae et al, 2019^21^. The three clusters in Dravnieks (block-rows demarcated by dashed blue lines) were obtained by permuting rows and columns independently using spectral bi-clustering. The rows of Chae were then permuted with the same row indices. Low variance features from both datasets were dropped for purposes of visual clarity.

### PYRFUME CODEFEST

On April 20^th^, 2022 we held the “Pyrfume Codefest” at the Association for Chemoreception Sciences (AChemS) annual conference. The goal was for attendees to learn and apply programming skills, collaborate with others, get feedback, explore others’ work, and connect with the larger AChemS community. The event was modeled after the Tidy Tuesday Project^29^. At an orientation event we provided two examples of starter code, reproducing a figure from Gilbert & Wysocki 1989^17^ and visualizing the Dravnieks Odor Atlas^16^. We oriented attendees to the available datasets, and provided a team of teachers to support attendees working in either R or Python. The goal was to welcome coders of all skill levels and from any chemosensory system.

We promoted the event with five Twitter threads spaced out in the week before the event. These tweets received over 6,500 impressions and 360 engagements. At the 2-hour event, 36 attendees put in 61 hours of coding and 116 hours overall. We received 18 code submissions at https://github.com/pyrfume/PyrfumeCodefest including a map of correct odor identifications in the National Geographic Smell Survey mapped by address; an exploration of the effect of allergies on odor identification; an exploration of the effect of age and sex on odor intensity and identification; and a shiny app (https://jmainland.shinyapps.io/PyrfumeDashboard/) that displays odor character for 7,262 molecules across seven datasets and provides search functionality to find a molecule with a particular odor.

## CONCLUSION

Pyrfume offers scientists, engineers, and trainees the opportunity to discover, test, and explore the world of experience through access to an unprecedented volume and diversity of data linked through a standardized format. Standardization allows for cross-modal analyses, meta-analyses, and benchmark construction for the next generation of predictive models. The next step is up to the broader research community, who we welcome to utilize this resource and, where applicable, to contribute their own datasets to increase their visibility and utility. We hope that this resource will bring to a close the “data-limited” era of olfaction research, and raise the bar for theoretical efforts.

## ACKNOWLEDGEMENTS

We thank all of those who contributed datasets to the project, the AChemS conference committee for allowing us to organize a Codefest, and NIH for support under R01DC018455. JBC was supported under NSF1553270.

## DATA AVAILABILITY

Pyrfume packages are available for python via pypi (`pip install pyrfume`) and R via CRAN (`install.packages(“rfume”)`). All datasets are available through the packages or directly on GitHub at http://github.com/pyrfume/pyrfume-data. Pyrfume is intended for non-commercial purposes under fair use principles, except where licenses permit commercial use. More information including full documentation and links to source code can be found at http://pyrfume.org.

